# Visualizing a protonated RNA state that modulates microRNA-21 maturation

**DOI:** 10.1101/852822

**Authors:** Jared T. Baisden, Joshua A. Boyer, Bo Zhao, Qi Zhang

**Affiliations:** Department of Biochemistry and Biophysics, University of North Carolina at Chapel Hill, Chapel Hill, North Carolina, USA; Department of Chemistry, University of North Carolina at Chapel Hill, Chapel Hill, North Carolina, USA

## Abstract

MicroRNAs are evolutionarily conserved small, non-coding RNAs that regulate diverse biological processes. Due to their essential regulatory roles, microRNA biogenesis is tightly regulated, where protein factors are often found to interact with specific primary and precursor microRNAs for regulation. Here, using NMR relaxation dispersion spectroscopy and mutagenesis, we reveal that the precursor of oncogenic microRNA-21 exists as a pH-dependent ensemble that spontaneously reshuffles the secondary structure of the entire apical stem-loop region, including the Dicer cleavage site. We show that the alternative excited conformation transiently sequesters the bulged adenine into a non-canonical protonated A^+^–G mismatch, conferring a two-fold enhancement in Dicer processing over its ground conformational state. These results indicate that microRNA maturation efficiency may be encoded in the intrinsic dynamic ensemble of primary and precursor microRNAs, providing potential means of regulating microRNA biogenesis in response to environmental and cellular stimuli.

---

MicroRNAs (miRNAs) are highly conserved, small noncoding RNAs that regulate more than 60% of protein coding genes at the post-transcriptional level^1–5^. Most miRNAs are initially transcribed by RNA polymerase II as introns of protein-coding genes or from independent coding genes into long primary transcripts (pri-miRNAs) that feature 5’-end 7-methylguanosine caps and 3’-end poly-A tails^6, 7^. In the canonical biogenesis pathway, pri-miRNAs are subsequently processed into ∼70 nucleotide precursor hairpins (pre-miRNAs) by the Microprocessor complex, consisting of one RNase III family enzyme, Drosha, and two DiGeorge critical region 8 proteins (DGCR8)^8, 9^. Pre-miRNAs are then exported from the nucleus to the cytoplasm by Exportin-5 (Ref. 10) and further processed into ∼20 base-pair miRNA/miRNA* duplexes by another RNase III family enzyme, Dicer, in complex with transactivation-responsive RNA binding protein (TRBP)^11, 12^. The resulting single-stranded mature miRNA is incorporated into the miRNA-inducing silencing complex (miRISC), which regulates protein expression by repressing translation, promoting deadenylation, and/or cleaving target mRNA^13^.

Due to their essential regulatory roles, miRNA biogenesis is tightly regulated to ensure proper gene expression^3–5^, and abnormal miRNA regulation has often been associated with cancer, neurological disorders, cardiovascular diseases and others^14, 15^. Remarkably, despite sharing the same set of enzymes in the canonical biogenesis pathway, individual miRNAs exhibit cell-type and cell-state specific expressions. Even those clustered on the same primary transcript can be differentially processed in a tissue-specific manner^3–5^. Over the past decade, it has been shown that specific sequences and structures of primary and precursor miRNAs can be recognized by processing machineries and protein factors for regulation^16–26^. For example, pri-miRNAs that possess a UGU motif in the apical loops are preferentially processed by the Microprocessor^19, 20^, pre-miRNAs encoding a two-nucleotide distance between the cleavage sites and the apical bulge/loop structures are more accurately processed by Dicer^21^, and miRNAs that feature stable basal stems in pri-miRNAs and flexible apical loops in pri-/pre-miRNAs are more efficiently processed by biogenesis machineries^22^. In addition, altering secondary structures and even primary sequences of pri-/pre-miRNAs via protein binding^23^, enzymatic-driven nucleotide modification^24, 25^ and disease-linked mutation^26^ can further influence the outcome of miRNA biogenesis^3^. During these regulatory processes, it is often perceived that protein factors act on the largely passive primary and precursor miRNAs to direct their maturation outcome. However, despite many non-coding RNAs having been shown to actively explore their conformational dynamics for function^27^, it remains elusive whether pri-/pre-miRNAs can play a role in modulating miRNA biogenesis in the absence of protein factors. This is largely due to our limited high-resolution structural and dynamic knowledge of most pri-/pre-miRNAs, where key regulatory elements of pri-/pre-miRNAs, such as the apical stem-loop region also known as the pre-element region, are often too flexible to be studied by conventional structural biology approaches.

Here, using NMR relaxation dispersion (RD) spectroscopy and Dicer processing assays, we set out to characterize the structural dynamics of microRNA-21 precursor (pre-miR-21) and examine how the intrinsic RNA conformational plasticity may contribute to miRNA maturation. MicroRNA-21, one of the first identified human miRNAs^28^, functions as an oncogene involved in tumorigenesis, progression, metastasis, and cell survival^29^, where its biogenesis is regulated at both transcriptional and post-transcriptional levels^30^. Previous studies have shown that the pre-element region of pri-/pre-miR-21 serves as an important element for regulating miR-21 biogenesis. Mutations that stabilize the pre-element inhibit Microprocessor processing of pri-miR-21 (Ref. 31), whereas binding of the KH-type splicing regulatory protein (KSRP) at this location promotes enzymatic processing of pri-/pre-miR-21s^32^. By carrying out NMR RD measurements, we discovered that the pre-element region of pri-/pre-miR-21 exists as a pH-dependent ensemble, which undergoes a two-state structural transition and dynamically accesses a low-populated (∼ 1 – 15%) transient, yet kinetically stable (lifetime ∼ 0.8 ms) state referred to as an excited state (ES) across physiologically relevant ranges of pH (pH ∼ 6.5 – 8.0). With ^15^N chemical exchange saturation transfer (CEST) NMR spectroscopy, we were able to directly measure, for the first time, an adenine N1 protonation event, which occurs at the Dicer cleavage site and underlies this unique, pH-dependent structural transition. This adenine protonation corresponds to a concerted secondary structural reshuffling of the entire pre-element region, transitioning the adenine from a bulged residue in the ground-state (GS) conformation to being sequestered into a non-canonical A^+^(*anti*)–G(*syn*) base pair in the ES. We further demonstrated that these distinct structures are processed differently by Dicer, where the ES-mimicking substrate is processed to mature miR-21 with a two-fold enhancement in efficiency over its GS counterpart. Hence, despite adopting an apparently simple secondary structure, pre-miR-21 encodes a dynamic ensemble at its pre-element region that encapsulates environmentally sensitive states with distinct fitness for processing. With the emerging view of RNA ES as a ‘hidden’ layer for regulation^27^, our results further suggest that miRNA processing intermediates may employ ES-encoded dynamic ensembles as potential means to regulate microRNA biogenesis in response to environmental and cellular stimuli.

## RESULTS

### The pre-element region of pre-miR-21 samples distinct conformational states

The miR-21 precursor consists of the pre-element region and miR-21/miR-21* helix, and is predicted to fold into a hairpin structure with four double-stranded helices, three bulges, and one apical loop (Fig. 1a). To focus on the pre-element region, we designed a shorter RNA construct, preE-miR-21, which contains the entire pre-element and the adjacent helix from the miR-21/miR-21* stem (Fig. 1b). NMR ^1^H-^1^H NOESY experiment on the imino region provides an excellent characterization of RNA secondary structure, as one imino resonance is expected for a canonical Watson-Crick base pair and two imino resonances are expected for a G-U wobble pair. Except for the formation of Watson-Crick base pairs at the lower stem, the pre-element of miR-21 does not adopt a stable conformation, as only weak imino resonances of G-U wobble pairs can be observed in the NMR ^1^H-^1^H NOESY spectrum (Fig. 1c). This observation is consistent with previous NMR studies on miR-21 precursor^33, 34^, where mutations of the pre-element were made to quench the structural flexibility into a single conformational state^33^. When we carried out an NMR ^13^C-^1^H HSQC experiment that probes non-solvent-exchangeable signals, surprisingly only 20 out of a total of 29 expected NMR resonances from preE-miR-21 were observed at room temperature (Fig. 1c). This spectroscopic behavior resembles the typical NMR phenomenon of exchange broadening, where interconversion between two or more states can lead to the disappearance of NMR signals. Indeed, by raising the temperature from 25°C to a more physiologically relevant 35°C, most of the missing resonances reappeared in the NMR ^13^C-^1^H HSQC spectrum (Fig. 1c), confirming the presence of conformational exchange.

**Figure 1.**
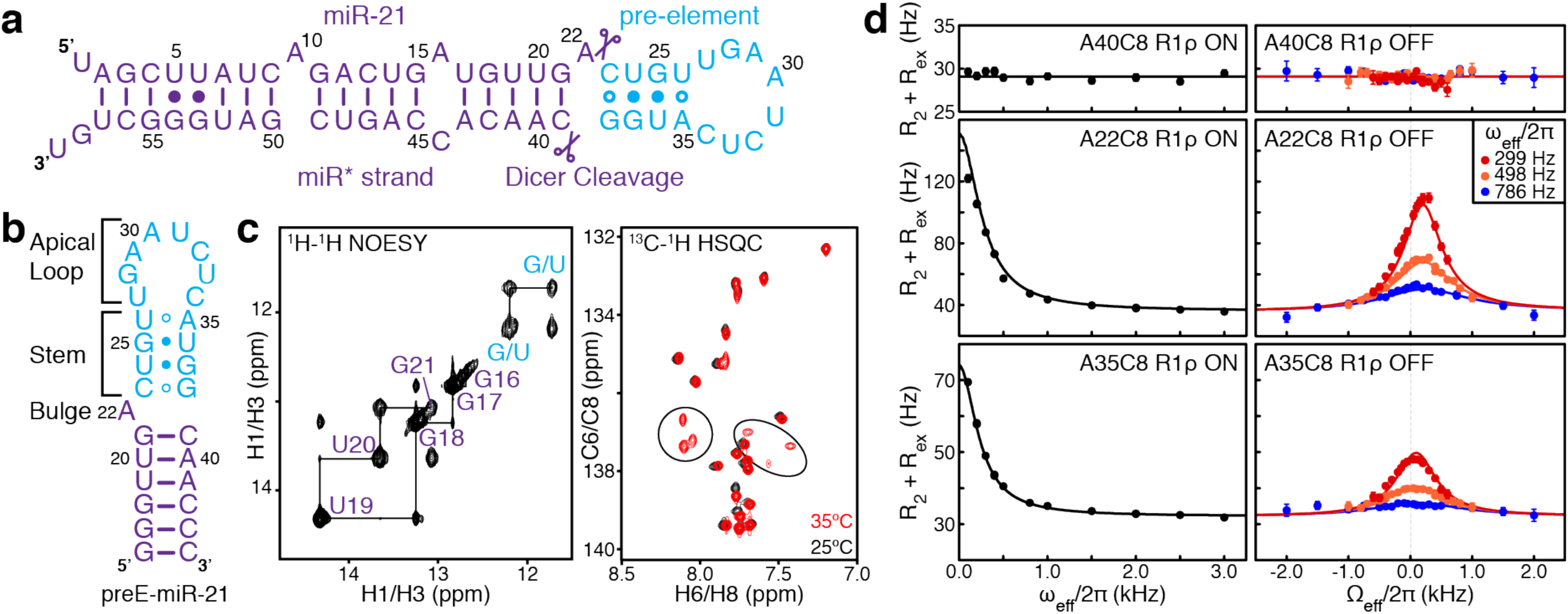
NMR characterization of preE-miR-21. (**a**) Secondary structure of pre-miR-21 with dicer cleavage sites highlighted as scissors. (**b**) Secondary structure of preE-miR-21 construct derived from NMR data, where Watson-Crick base pairs, GU wobbles, and potential Watson-Crick base pairs are highlighted with lines, filled circles, and open circles, respectively. (**c**) NMR ^1^H-^1^H NOESY spectrum of the imino proton region of preE-miR-21 at 10°C and ^13^C-^1^H HSQC spectra of base carbon (C6 and C8) region of preE-miR-21 at 25°C and 35°C. (**d**) Representative ^13^C on-resonance and off-resonance relaxation dispersion (RD) profiles at 35°C showing dependence of *R*_2_ + *R*_ex_ on spin-lock power (ω_eff_/2π) and offset (Ω/2π), respectively, where Ω is the difference between the spin-lock carrier frequency and the observed resonance frequency. RD profiles of A40 are fit to a single-state model and RD profiles of A22 and A35 are fit to a global two-state model using the Bloch-McConnell equation. Error bars are experimental uncertainties (s.d.) estimated from mono-exponential fitting of *n* = 3 independently measured peak intensities.

Recent developments of NMR R_1ρ_ RD spectroscopy have opened new avenues to quantify microsecond-to-millisecond conformational changes and made it possible to study RNA ESs that are too low-populated and short-lived to be detected by conventional techniques^35–39^. Here, we carried out both on-resonance and off-resonance low spin-lock field R_1ρ_ RD experiments to quantify the exchange process in preE-miR-21. For residues from the miR-21/miR-21* stem region, we observed flat RD profiles for base (C2, C5, C6, and C8) and sugar (C1’) carbons (Fig.1d and Supplementary Fig. 1), which are consistent with one stable helical conformation of this region. In contrast, residues within the pre-element region, ranging from the bulge residue A22 to the stem residue A35 (Fig. 1d and Supplementary Fig. 2), display power and offset dependent RD profiles. Indeed, these RD profiles can be global-fitted to a single two-state (GS ↔ ES) exchange process. These results reveal that the pre-element region is not only conformationally flexible, but also dynamically interconverts between at least two structurally and kinetically distinct states, where the ES has a low population (*p*_ES_) of 15.2 ± 0.3% and a short lifetime (τ_ES_ = 1/*k*_EG_) of 816 ± 15 μs (Fig. 1d).

### The ES involves transient protonation at the Dicer cleavage site

To gain structural insights into the ES, we utilized NMR chemical shifts, which are one of the most sensitive measurements for probing local chemical environments. We examined ES carbon chemical shifts 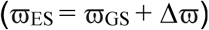, where 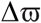 is the chemical shift difference between ES 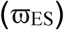 and GS 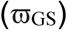 extracted from the two-state analysis of an R_1ρ_ RD profile (Supplementary Fig. 3). Among all the extracted ES chemical shifts, the base carbon C8 of bulge A22, which resides at the Dicer cleavage site, displays the largest deviation 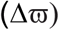 with a ∼1.49 ppm down-field shift from its GS position (Fig. 1d and Supplementary Fig. 3). Notably, we were not able to obtain the RD profile for base carbon C2 of A22, as the C2H2 resonance remains severely broadened beyond detection in the NMR ^13^C-^1^H HSQC (Fig. 2a), suggesting even larger perturbations in carbon C2 and/or proton H2 chemical shifts between ES and GS. The dramatically different behavior of C8H8 and C2H2 resonances from the same base is reminiscent of recent NMR studies on transiently N1-protonated adenines^40, 41^.

**Figure 2.**
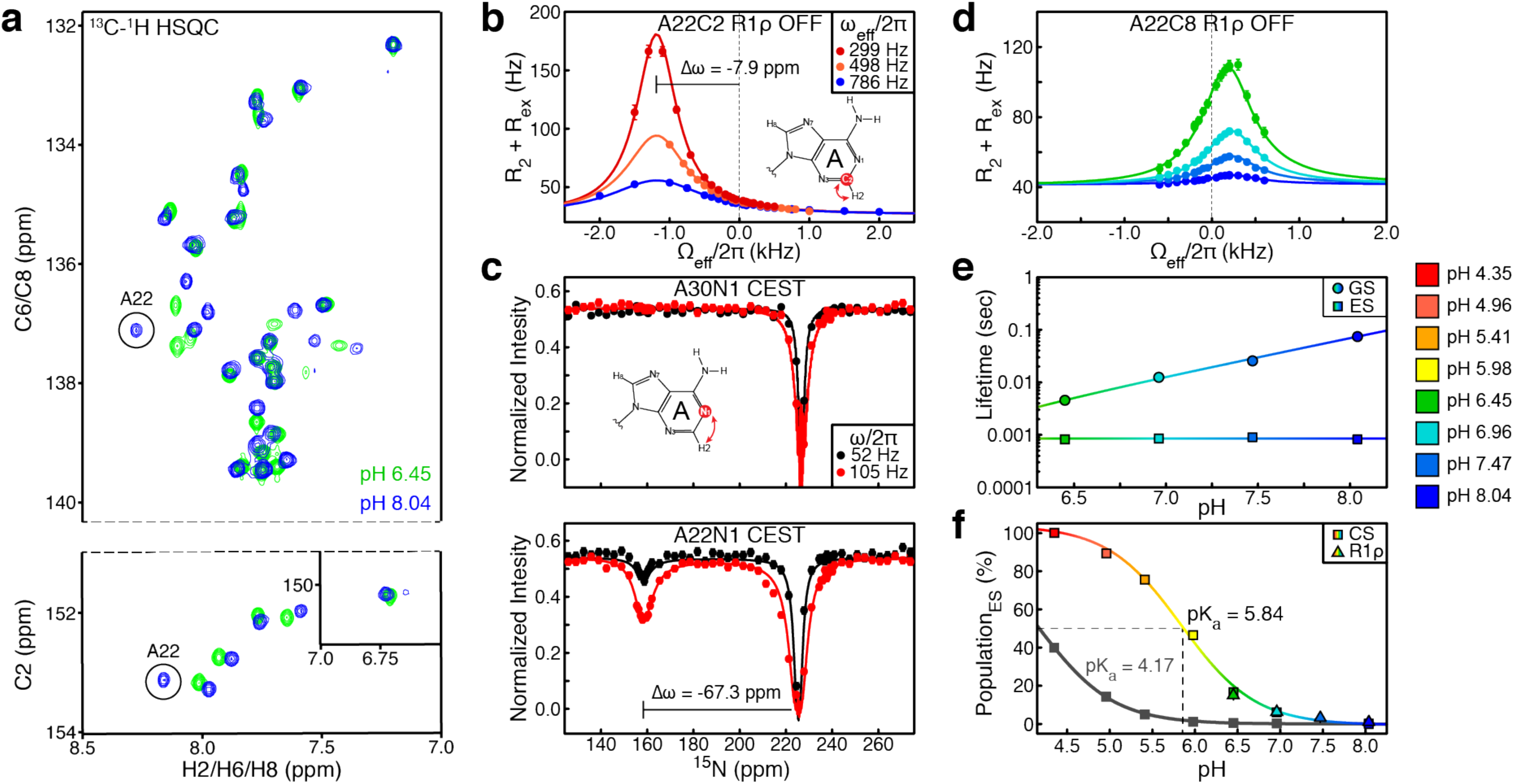
PreE-miR-21 populates a protonated excited state with a neutral shifted p*K*_a_. (**a**) NMR ^13^C-^1^H HSQC spectra of base carbon (C2, C6, and C8) region of preE-miR-21 at pH 6.45 and pH 8.04. (**b**) ^13^C off-resonance RD profiles of A22-C2 at pH 8.04. (**c**) ^15^N CEST profiles of A30-N1 and A22-N1 at pH 8.04, which are fit to a single-state and a two-state model, respectively, using the Bloch-McConnell equation. (**d**) The pH-dependent ^13^C off-resonance RD profiles of A35-C8 at spin-lock power of ω_eff_/2π =299 Hz. (**e**) The pH-dependent apparent lifetimes of GS and ES from RD analysis. (**f**) The pH-dependent population of the excited state based on R_1ρ_ RD data and A22-C8 chemical shift (CS) for extracting an apparent pK_a_ of A22. Representative pK_a_ derived from unpaired A29, A30 and A35 is shown in black. Error bars are experimental uncertainties (s.d.) estimated from mono-exponential fitting of *n* = 3 independently measured peak intensities.

To examine whether the exchange process could be due to possible protonation events, we increased the pH of the sample from 6.45 to 8.04, aiming to shift the equilibrium towards non-protonated states. Indeed, the NMR ^13^C-^1^H HSQC spectrum recorded at pH 8.04 exhibits much higher quality, where exchange broadening of most resonances is substantially reduced, such that the A22-C2H2 resonance can be readily observed (Fig. 2a). Low spin-lock field NMR R_1ρ_ RD measurements provide further quantitative support that the observed exchange involves a protonated ES of the pre-element region of miR-21 (Fig. 2b and Supplementary Fig. 4). Global fit of RD profiles showed that the GS ↔ ES equilibrium is significantly shifted towards GS at pH 8.04, where the ES population (*p*_ES_) is reduced to a mere 1.1 ± 0.1%. In addition, a two-state analysis of the RD profile of base carbon A22-C2 further revealed a remarkable 7.9 ppm difference between its GS and ES chemical shifts, resulting in an ES chemical shift of 144.5 ppm (Fig. 2b and Supplemental Fig. 5). This significantly up-field shifted C2 chemical shift is consistent with C2 chemical shifts reported for stably N1-protonated adenines, strongly suggesting that A22 is protonated at the N1 site in the ES.

Recently, we have developed nucleic-acid-specific ^15^N CEST NMR spectroscopy to study RNA conformational exchanges using non-proton-bonded nitrogens as probes^42^. This technique also enables direct evaluation of the protonation status of adenines in low-populated and short-lived states, which have remained elusive to date. By measuring ^15^N CEST profiles at pH 8.04, we were able to unambiguously identify that A22 is transiently protonated at N1 (Fig. 2c). Unlike N1 of A30, which is not protonated and displays an apparent single-dip CEST profile, the nitrogen CEST profile of A22-N1 exhibits two distinct intensity dips that correspond to two alternative conformations. A two-state analysis of the CEST profile validates that A22-N1 probes the same two-state exchange process, where extracted ES population (*p*_ES-CEST_ ∼ 1.1 ± 0.1%) and lifetime (τ_ES-CEST_ ∼ 645 ± 114 μs) from ^15^N CEST are very similar to those obtained from ^13^C R_1ρ_ RD at pH 8.04 (*p*_ES-R1ρ_ ∼ 1.1 ± 0.1%, τ_ES-R1ρ_ ∼ 823 ± 109 μs). The ES chemical shift of A22-N1 directly supports a protonated N1 with an unprecedented up-field shift of 67.3 ± 0.2 ppm to 157.7 ppm, residing well among resonances of proton-bonded imino nitrogens in RNA (∼134-152 ppm in Gs and ∼154-165 ppm in Us).

To obtain more insights into the A22 (GS) ↔ A22^+^ (ES) transition, we further carried out carbon R_1ρ_ RD and nitrogen CEST measurements on base (C2, C8, N1) and sugar (C1’) moieties of A22 at pH 6.96 and 7.47 (Fig. 2d and Supplementary Fig. 5). Consistent with being a protonation-dependent process, the population of A22^+^ gradually increases from ∼ 1% at pH 8.04 to ∼ 15% at pH 6.45. Surprisingly, the lifetime of the ES A22^+^ remains largely unperturbed between pH 6.45 and pH 8.04, where the average lifetime is τ_ES_ ∼ 847 ± 49 μs (Fig. 2e). In contrast, the apparent lifetime of the GS, which is derived from the extracted rate of exchange (τ_GS_ = 1/*k*_GE_), reduces substantially from τ_GS_ ∼ 74 ms at pH 8.04 to τ_GS_ ∼ 5 ms at pH 6.45 (Fig. 2e). The high population of A22^+^ at pH 6.45, which otherwise would be close to zero based on the intrinsic pK_a_ (∼ 3.5) of free adenine N1 site^43^, further suggests that A22 has a distinct protonation propensity when compared with other unstructured adenines. Consistent with this observation, pH-dependent chemical shift analyses showed a pK_a_ value of 5.84 ± 0.08 for A22, which is substantially shifted towards neutral pH from adenines in the apical-loop that have an average pK_a_ value of 4.17 ± 0.06 (Fig. 2f and Supplementary Fig. 6). Taken together, these results unambiguously revealed that preE-miR-21 undergoes a pH-dependent conformational transition, where A22 at the Dicer cleavage site is specifically protonated in the ES.

### The transient protonation couples global secondary structural reshuffling

How does the excited state stabilize a site-specific protonation? To address this, we first evaluated the role of each structural motif of the pre-element region – the bulge, the stem, and the apical loop – in the observed conformational transition (Fig. 3a-c and Supplementary Fig. 7). Bulge A22 is the site of protonation. Without A22, we could not detect any conformational exchange within the rest of the pre-element region, as evidenced with flat RD profiles for the A22-deletion mutant (Fig. 3a). Not only is a protonated A22 the result of the structural transition, this protonation may likely be the chemical basis that triggers the larger transition across the entire pre-element region. In addition to the indispensable bulge A22, we found that both a weak stem and a flexible apical loop are needed to achieve the structural transition. Stabilizing the two G-U wobble pairs with G-C Watson-Crick pairs completely quenches the exchange (Fig. 3b); replacing the apical loop with a highly structured UUCG tetraloop also eliminates the transition (Fig. 3c). These results suggest that the pre-element region serves as a unified structural entity to enable a concerted transition towards stabilizing the protonated excited state. The rate of exchange (*k*_ex_ = *k*_GE_ + *k*_EG_ ∼ 1445 s^-1^) is an order of magnitude slower than rates observed from local structural changes involving transient adenine protonation^44^, but similar to the secondary-structure-based long-range communication observed HIV-1 TAR RNA^45^, further supporting a global secondary structural reshuffling of the pre-element region.

**Figure 3.**
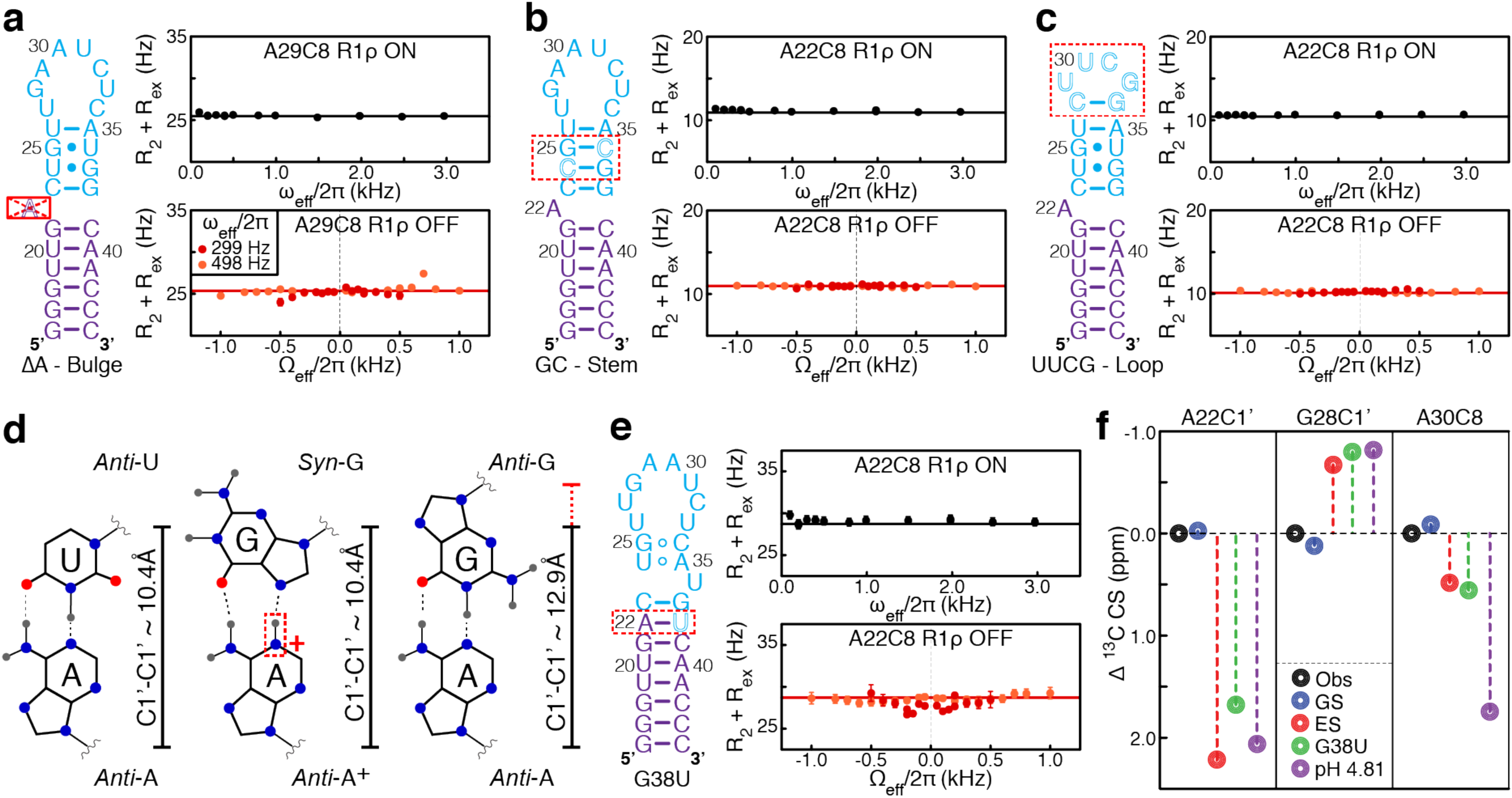
Excited-state structure of preE-miR-21. (**a-c**) Secondary structures and representative ^13^C on-resonance and off-resonance RD profiles of bulge, stem, and loop mutants. (**d**) Sugar-sugar (C1’-C1’) distances of A-U Watson-Crick base pair, A-G mismatch, and A^+^-G mismatch. (**e**) Secondary structure and representative ^13^C on-resonance and off-resonance RD profiles of ES-mimic mutant. (**f**) Comparison of carbon chemical shifts for the GS, ES, the mutant mimics, and wild-type construct at pH 4.81. Error bars are experimental uncertainties (s.d.) estimated from mono-exponential fitting of *n* = 3 independently measured peak intensities.

To further delineate the secondary structure of the excited state, we employed a mutate-and-chemical-shift-fingerprinting strategy^44^. In this approach, mutations are introduced to stabilize conformational features unique to a proposed/predicted ES secondary structure, which are then validated by comparing chemical shift differences between the mutant and wild-type 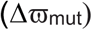 to those extracted from NMR RD profiles 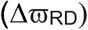. Here, we used MC-fold^46^ to predict possible alternative low-energy secondary structures of preE-miR-21. Strikingly, most of the predicted structures share a common feature of an A22-G38 base pair, whereas the remaining pre-element adopts various secondary structures that are distinct from the ground-state conformation (Supplementary Fig. 8). Being protonated at N1, A22 could potentially be base paired with G38 in the ES, albeit in the A^+^(*anti*)–G(*syn*) form rather than the conventional A(*anti*)–G(*anti*) pair, where the *syn* conformation of G38 is indicated with the down-field chemical shift of base carbon C8 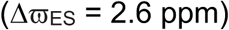 at pH 8.04 (Supplementary Fig. 4). An interesting structural feature of the A^+^(*anti*)–G(*syn*) pair is that it largely retains an overall A-form-like geometry with an inter-sugar distance of 10.4 Å^47^, whereas the A(*anti*)–G(*anti*) pair substantially widens this distance to 12.9 Å^48^ and subsequently distorts the helical geometry of neighboring base pairs (Fig. 3d).

To test this proposed ES structural feature, we mutated G38 to a uridine, which not only sequesters A22 into a base pair, but also maintains an A-form geometry at the site of mutation. The G38U mutant converges into a single state as evidenced by flat RD profiles (Fig. 3e and Supplementary Fig. 9), and largely represents the ES of preE-miR-21 based on chemical shifts. Good agreement was observed between 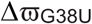 and 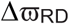 for 15 out of 19 base and sugar carbon resonances from the pre-element residues with detectable RD profiles, including A22(C1’), C23(C1’), G28(C1’), A29(C2/C8), A30(C8), U31(C1’), C32(C1’), U33(C6), C34(C6), A35(C1’/C2/C8), U36(C6), and C39(C5) (Fig. 3f and Supplementary Figs. 3 and 4). The agreement of sugar carbon C1’s of A22 and C23 further supports the ES adopting an A-form-like backbone geometry at the site of protonation. The deviations between 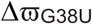 and 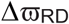 for base carbons C2 and C8 of A22 can be attributed to protonation-induced major chemical shift perturbations in the wild-type, which cannot be recapitulated with this mutation. However, the deviations for C32-C5 and U36-C1’ could be due to their relatively small chemical shift differences (<0.5 ppm) and/or local conformational perturbations in the wild-type from the mutation, which is subject to future investigation. As the G38U mutant closely mimics the ES, it also provides some insights into the two residues (U24 and U26) in the pre-element region that showed no detectable RD. For U24-C6, its flat RD profile can be explained with essentially identical chemical shifts between the ground and excited states as indicated with 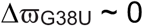, whereas the lack of detectable RD for U26-C6 may be due to additional local conformational perturbations that are subject to further studies (Supplementary Fig. 9). To provide independent validation of the chemical shift fingerprints of the ES, we compared 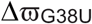 to chemical shift differences of the wild type between pH 8.04 and pH 4.81 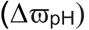, where the low pH value was chosen to shift the population towards the protonated ES without inducing global protonation of adenines and cytosines. Good agreement was observed between 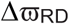 and 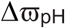, except for some major deviations from unpaired adenines and cytosines that are likely due to rapid protonation at pH 4.81 given their intrinsic pK_a_s (∼ 3.5 – 4.2)^43^ when unpaired (Fig. 3f and Supplementary Figs. 3 and 4). In particular, excellent agreement between 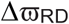 and 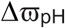 of A22-C2 and A22-C8 complements the G38U mutant. Taken together, low pH and G38U are each able to recapitulate a portion of the ES structure, with low pH chemical shifts matching changes in A22 residues, and G38U chemical shifts matching throughout the rest of the structure. These results strongly suggest that pre-miR-21 undergoes a global structural reshuffling at the pre-element region, where A22 is transiently protonated and forms a distinct A^+^(*anti*)–G(*syn*) base pair in the ES.

**Figure 4.**
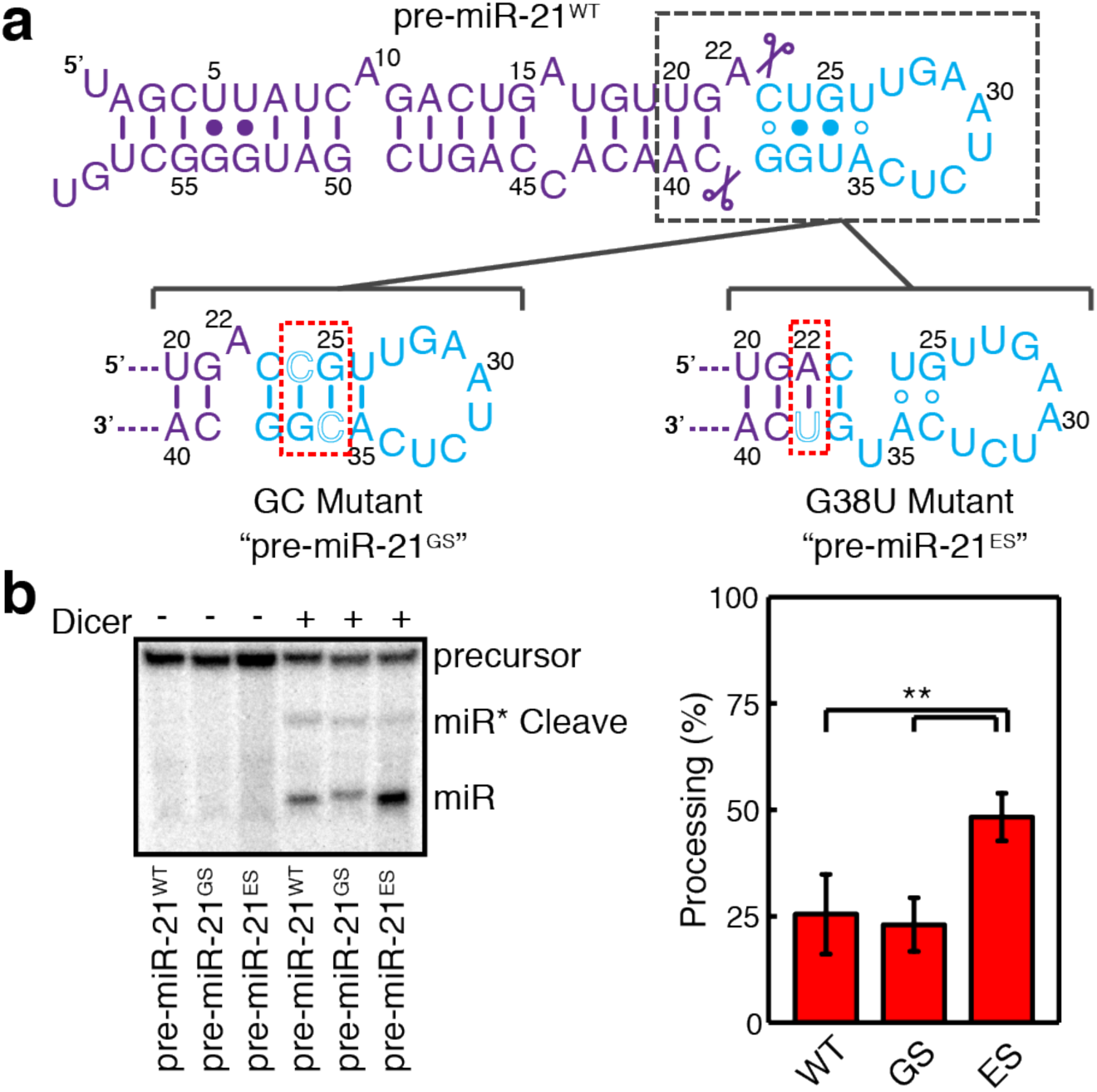
Excited state of preE-miR-21 enhances dicer processing. (**a**) Secondary structures of the wild-type, ground-state mimic, and excited-state mimic of pre-miR-21. (**b**) Dicer processing assays of wild-type and mutant pre-miR-21s, quantification using ImageQuant shown on right (** denotes p ≤ 0.01 via student T-Test). Shown are means and standard deviations (s.d.) from *n* = 4 independent assays.

### The miR-21 precursor encodes states with differential Dicer processivities

The conformational transition of pre-miR-21 is the protonation-driven base pairing of the bulged A22 that resides specifically at the location of Dicer cleavage. Hence, it is of interest to see how these structural changes may affect Dicer cleavage of the miR-21 precursor. To examine this, we designed GS- and ES-mimicking substrates and performed processing assays using commercially available recombinant human Dicer (Fig. 4a). For the GS-mimicking substrate (pre-miR-21^GS^), we mutated the two G– U wobbles with two G–C pairs, which was shown to stabilize the flexible stem that forms in the ground state. For the ES-mimicking structure (pre-miR-21^ES^), we incorporated a G38U mutation to the full-length pre-miR-21. We would like to note that the G38U mutant, which recapitulates key ES structural features, cannot perfectly mimic the electrostatic property of the protonated ES. However, since Dicer cleaves the phosphate backbone, we anticipate structures of the backbone, rather than the base pairing identity of the ES, may influence Dicer activity. In order to generate a native-like precursor with 5’-terminal phosphate group and sequence, we fused a hammerhead ribozyme to the 5’-end of full-length miR-21 precursor. During *in vitro* transcription, the hammerhead ribozyme self-cleaves, and the resulting miR-21 precursor is 5’-end phosphorylated and labelled with [γ-^32^P] ATP. Remarkably, GS- and ES-mimicking substrates exhibited substantially different Dicer processivities (Fig. 4b). As can be seen, ∼ 23 ± 6% of pre-miR-21^GS^ was cleaved by Dicer to generate mature miR-21, which is essentially identical to a processivity of ∼ 26 ± 9% for the wild-type substrate (pre-miR-21^WT^). This is consistent with pre-miR-21^WT^ occupying ∼99% GS under the assay condition of pH 8.04. In contrast, Dicer processed pre-miR-21^ES^ much more efficiently than its GS counterpart, where double the amount of substrate (∼ 48 ± 6%) was converted to mature miR-21. Together, these results not only unveil differential fitness of the GS and ES of the pre-element region of the pre-miR-21 for miR-21 maturation, but further exemplify the importance of RNA structures in directing the overall outcome of miRNA biogenesis.

## DISCUSSION

Here, by integrating structural, dynamic, and functional analyses on miR-21 precursor, we showed that the intrinsic conformational plasticity of miRNA processing intermediates can serve as a new layer of regulation for miRNA biogenesis. Often, structural changes in pri-/pre-miRNAs can be induced upon binding to protein regulators^23^, nucleotide modifications such as ADAR1-mediated adenine-to-inosine editing^24^ and METTL1-mediated methylation^25^, and disease-linked mutations^26^. RNA structural motifs are also important factors in recognition by processing machineries for miRNA biogenesis, where altering primary and/or precursor structures of a target miRNA can further lead to altered biogenesis, inducing a change in physiological outcomes^3^. In contrast to these adaptive changes, we found that pre-miR-21 encodes a dynamic ensemble in its apical stem-loop region that undergoes spontaneous structural transitions between two kinetically and functionally distinct states (Fig. 5). In the ground state, the two Dicer cleavage sites reside within a largely unstructured region; in contrast, both locations become structured in the excited state. Relative to the ground state, the excited state conformation more closely resembles the optimal structure for Dicer, where both cleavage sites are base-paired and positioned two-nucleotides away from a flexible apical loop^21^, hence, providing a better topology for Dicer cleavage.

**Figure 5.**
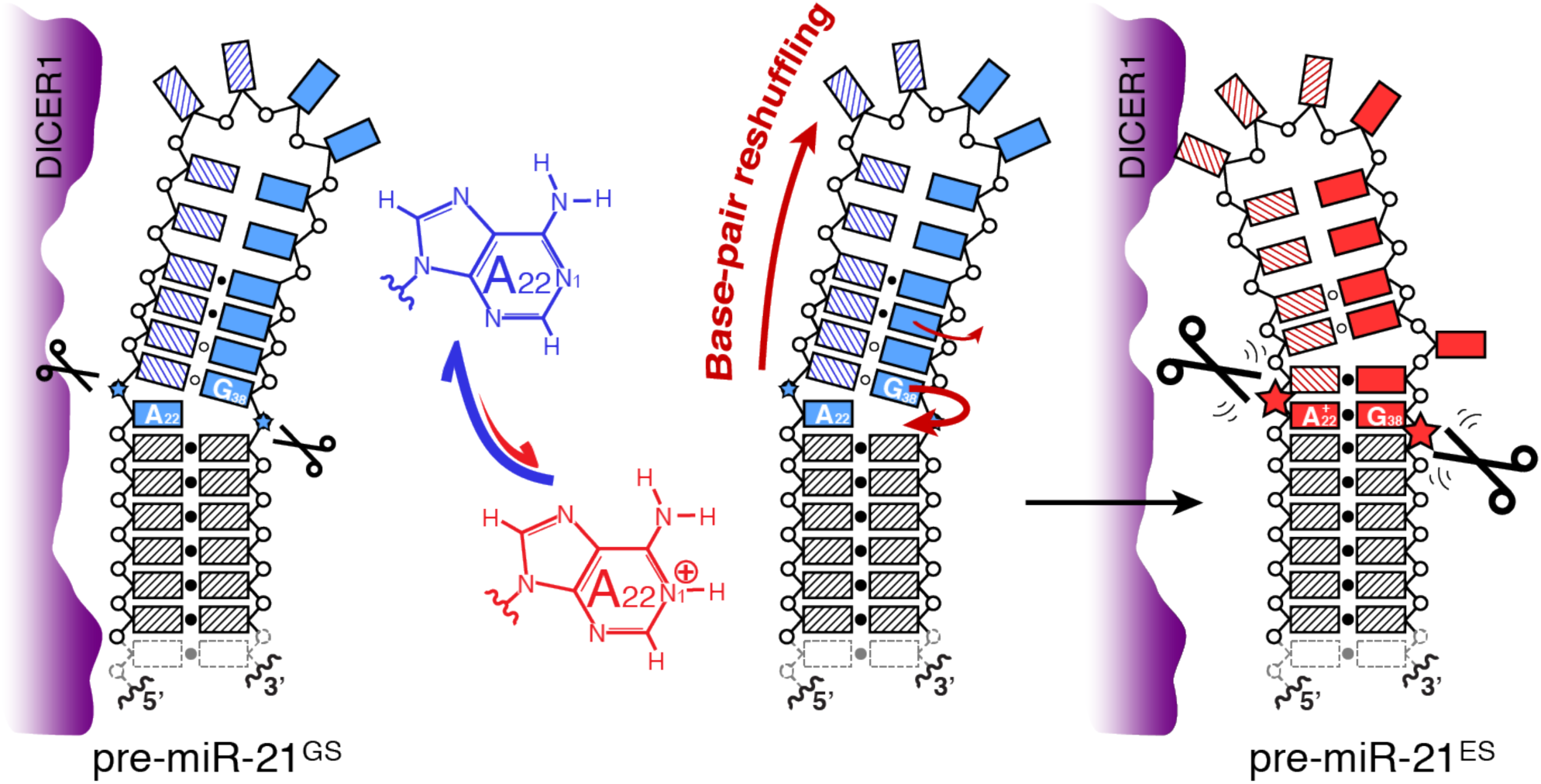
Modulation of miR-21 maturation with a protonation-dependent structural ensemble. Protonation at N1 site of A22 sequesters the bulged adenine into a non-canonical A^+^–G mismatch and is associated with a long-range conformational reshuffling of the pre-element region. This structural rearrangement results in a conformation that is better suited for Dicer processing to generate mature miR-21.

A hallmark of the ES of pre-miR-21 is protonation of the adenine residue at the Dicer cleavage site. Protonation is a fundamental chemical property and one of the smallest chemical modifications on nucleic acids^43^. The intrinsic pK_a_s for protonation of adenines and cytosines are acidic and reside far from the physiological pH ranges. However, by adopting sophisticated structures, RNA can shift acidic pK_a_s toward neutral pHs, such that specific ionization can be achieved under physiological conditions for function^43^. For example, the universally conserved adenine residue at the active site of the ribosome has a shifted pK_a_ to serve as a general acid-base catalyst for peptide formation^49^, whereas the murine leukemia virus (MLV) recoding signal employs a protonated adenine as a structural factor to stabilize a compact pseudoknot in order to allow the ribosome to bypass the Gag stop codon^40^. Here, protonation provides the crucial chemical basis for pre-miR-21 to form the A^+^(*anti*)–G(*syn*) base pair, which ensures the cleavage site adopts an overall A-form-like topology in the ES. Without being protonated, the adenine residue may still be able to pair with the upper stem guanine residue, albeit in the form of A(*anti*)–G(*anti*) mismatch, which is likely functionally indistinguishable from the GS. Another feature of the protonation event in pre-miR-21 is that the underlying structural transition occurs at the millisecond timescale, which is substantially faster than those involved in major structural changes, such as the adenine protonation in MLV^40^. This fast GS ↔ ES interconversion could enable pre-miR-21 to rapidly reach new equilibrium upon a transient high acid load due to disease-induced metabolic shifts, modulating maturation of miR-21 in response to environmental stimuli. Despite displaying distinct *in vitro* outcomes, a functional understanding of the role of pre-miR-21 protonation in regulating biogenesis will require future investigations that evaluate the response of pre-miR-21 under various cellular conditions such as hypoxia and acidosis.

Interestingly, the spontaneous conformational transition in pre-miR-21, which involves secondary structural reshuffling of the pre-element region, is reminiscent of that observed in Lin28-dependent regulation of the biogenesis of let-7 family of miRNAs^50^. In general, RNA secondary structural changes encounter large kinetic barriers, hence, need to be catalyzed by external factors such as RNA-binding proteins. Both domains of Lin28 work cooperatively to bind two independent RNA elements to induce the regulatory structural changes of pri-/pre-let-7s. In contrast, pre-miR-21 accomplishes such structural changes without protein factors by accessing an excited state, where the bulged adenine base pairs with the upper stem guanine residue, propagating a global change in strand register. Here, all three structural elements of the apical stem-loop – the bulge, metastable stem, and flexible loop – are essential to achieve this concerted movement, and eliminating any of them abolishes the spontaneous transitions in pre-miR-21. While our observation represents the first example of an excited state in pre-miRNAs, an ES-based mechanism for remodeling distant motifs has also been recently reported in HIV-1 TAR RNA^45^. Since the apical stem-loop is a common structural feature among all pri-/pre-miRNAs, we speculate that many miRNA processing intermediates may encode similar ES-based conformational plasticity for long-range communication across their regulatory pre-element regions, which is further supported by a recent computational modeling of secondary structural ensembles of miRNAs^26^.

It has become increasingly clear that many non-coding RNAs (ncRNAs) do not fold into single static structures, instead, they dynamically interconvert between different conformational states for function^27^. Recent developments in NMR techniques have opened new avenues to probe RNA structural transitions involving alternative conformational states that often evade detection from conventional biophysical and biochemical methods due to their low populations and/or transient lifetimes^38, 39, 44^. Indeed, these technical advances have unveiled the presence of a diverse set of excited states in non-coding RNAs. Our discovery of the transiently protonated state in pre-miR-21 was also made possible by these new techniques. In particular, the development of a nucleic-acid-optimized ^15^N NMR CEST method has enabled, for the first time, direct identification of transient protonation in nucleic acids, illuminating the crucial chemical basis for delineating the relationship between structure and function. With the emerging view of ESs as a ‘hidden’ layer of regulation^27^, the growing repertoire of functional RNA ESs with distinct structural features promise novel strategies and developments in RNA-targeted therapeutics.

## METHODS

Methods, including statements of data availability and references, are available in the online version of the paper.

## Acknowledgements

We thank G. Young and S. Parnham for maintenance of NMR instruments and members of the Zhang lab for critical comments. This work was supported by start-up fund from the University of North Carolina at Chapel Hill and an NSF grant (MCB1652676).

## Author contributions

J.T.B. and Q.Z. conceived the project and experimental design. J.T.B., J.A.B., B.Z. and Q.Z. prepared the samples, carried out NMR experiments, analyzed the data, and wrote the paper.

## Additional information

Any supplementary information, chemical compound information and source data are available in the online version of the paper. Reprints and permissions information is available online at http://www.nature.com/reprints/index.html. Correspondence and requests for materials should be addressed to Q.Z.

