## Supplementary Information for "Visualizing a protonated RNA state that modulates microRNA-21 maturation"

### ONLINE METHODS

**Sample preparation.** Unlabeled, uniformly $^{13}C/^{15}N$-labeled, and adenine-specifically $^{13}C/^{15}N$-labeled preE-miR-21 samples and mutants were prepared as previously described[51]. Briefly, after *in vitro* transcription, samples were ethanol precipitated, gel purified using 15% denaturing polyacrylamide gel, electro-eluted with the Elutrap system (Whatman), purified with an 5 mL Hi-Trap Q anion-exchange column (GE Healthcare), and desalted by exchanging into $H_2O$ using an Amicon filtration unit with 3K Da MW cut-off membrane (Millipore). Samples were then concentrated and exchanged into NMR buffers with 50 mM KCl and 50 $\mu$M EDTA, where 10 mM acetate buffers were used for pHs at 4.35, 4.96, and 5.41, and 10 mM sodium phosphate buffers were used for pHs at 5.98, 6.45, 6.96, 7.47, and 8.04. For $H_2O$ samples, 5% $D_2O$ was added. For $D_2O$ samples in sodium phosphate buffer, $H_2O$ samples were lyophilized and redissolved in the same volume of 99.996% $D_2O$ (Sigma). For $D_2O$ samples in sodium acetate buffer, samples were exchanged into 10 mM acetate buffer with 50 mM KCl and 50 $\mu$M EDTA in $D_2O$.

**NMR spectroscopy.** All NMR experiments were carried out on a Bruker Avance III 600 MHz spectrometer equipped with a 5-mm triple-resonance (TCI) cryogenic probe. Exchangeable proton spectra were recorded using $H_2O$ samples at 283 K, and nonexchangeable proton spectra were recorded at 298 K and 308 K using $H_2O$ and $D_2O$ samples. Spectra were processed and analyzed with TOPSPIN 3.5 (Bruker), NMRPipe[52], NMRView[53], and Sparky 3.110 (University of California, San Francisco, CA). The assignments for preE-miR-21 were obtained with samples at pHs 6.45 and 8.04 and the assignments for its mutants were obtained with samples at pH 6.45 using 2D NOESY, 2D

TOCSY, $^1H$-$^{15}N$ HSQC, $^1H$-$^{13}C$ HSQC, 2D HCCH-COSY, and HCN experiments on unlabeled, uniformly labeled and adenine-specifically $^{13}C$/$^{15}N$ labeled samples with standard protocols[54]. The apparent $pK_a$ values of adenines were obtained by fitting pH-dependent excited-state population and chemical shift to the Henderson-Hasselbalch equation as described previously[55].

**$^{13}$C $R_{1\rho}$ relaxation dispersion measurements and data analysis.** The on- and off-resonance relaxation dispersion profiles were measured using the 1D selective $R_{1\rho}$ experiment developed by Al-Hashimi and co-workers[37] and a constant-time approach described by Kay and co-workers[36], where $R_{1\rho}$ values were obtained from a single delay period ($T_{EX}$). For on-resonance experiments, the relaxation delay was set to $T_{EX}$ = 32 ms; for off-resonance experiments, the relaxation delay was set to $T_{EX}$ = 40 ms, except $T_{EX}$ = 16ms for A22-C8 and A35-C1' and $T_{EX}$ = 24 ms for C23-C6, U36-C6, and G28-C1'. Relaxation rates were determined by fitting peak intensity to a single exponential decay as $R_{1\rho} = -\ln(I_T/I_0)/T_{EX}$ , where $I_T$ is the decayed peak intensity and $I_0$ is the reference peak intensity. Relaxation rate errors were estimated by intensity deviations between three duplicates at $T_{EX}$ = 0 and the signal-to-noise ratios in 1D spectra. The largest of the two errors was reported. For on-resonance experiments, eleven $^{13}$C spin-lock fields (ω/2π) of 100, 199 (x2), 299, 399, 498 (x2), 786, 982, 1474, 1965 (x2), 2456, and 2947 Hz were employed, (x2) indicates performed in duplicates. Due to large C-C couplings, the lowest $^{13}$C spin-lock field (ω/2π) of 100 Hz was not used in measuring on-resonance C5/C6/C1' RD profiles. For off-resonance experiments, three $^{13}$C spin-lock fields (ω/2π) of 299, 498, and 786 Hz were used. For ω/2π = 299 Hz, the $^{13}$C offsets ranged between -600 and 600 Hz with a spacing of 100 Hz and a smaller spacing of 50 Hz between -200 and 200 Hz;

for $\omega/2\pi$ = 498 Hz, the $^{13}C$ offsets ranged between -1000 and 1000 Hz with a spacing of 200 Hz from -1000 to -800 Hz and from 800 to 1000 Hz, a spacing of 100 Hz from -800 to -100 Hz and from 100 to 800 Hz, and single points at -50 and 50 Hz; for $\omega/2\pi$ = 786 Hz, the $^{13}C$ offsets ranged between -2000 and 2000 Hz with a spacing of 500 Hz from -2000 to -1000 Hz and 1000 to 2000 Hz, a spacing of 250 Hz from -1000 to -500 Hz and 500 to 1000 Hz, and a spacing of 100 Hz from -500 to -100 Hz and from 100 to 500 Hz, and single points at -50 and 50 Hz. $^{13}C$ spin-lock powers were calibrated according to the 1D approach by Guenneugues *et al.*[56] as previously described[51,57].

Relaxation dispersion profiles were analyzed as described previously[51]. Briefly, on- and off-resonance relaxation dispersion profiles were obtained by measuring the rate of decay of magnetization over the spin-lock period as a function of spin-lock power ($\omega_{eff}/2\pi$) and spin-lock offset ($\Omega$), respectively, where $\Omega = \omega_{rf} - \Omega_{obs}$ is the frequency difference between the spin-lock carrier frequency ($\omega_{rf}$) and the observed resonance frequency ($\Omega_{obs}$). The $R_{1\rho}$ profiles of residues displaying conformational exchange were fit to a two-state exchange model between the ground (G) and excited (E) states based on the Bloch-McConnell equation[58],

$$\frac{d}{dt}\begin{pmatrix} I_x^G \\ I_y^G \\ I_z^G \\ I_x^E \\ I_y^E \\ I_z^E \end{pmatrix} = \begin{pmatrix} -R_2^G-k_{GE} & -\omega_G & 0 & k_{EG} & 0 & 0 \\ \omega_G & -R_2^G-k_{GE} & -\omega_1 & 0 & k_{EG} & 0 \\ 0 & \omega_1 & -R_1^G-k_{GE} & 0 & 0 & k_{EG} \\ k_{GE} & 0 & 0 & -R_2^E-k_{EG} & -\omega_E & 0 \\ 0 & k_{GE} & 0 & \omega_E & -R_2^E-k_{EG} & -\omega_1 \\ 0 & 0 & k_{GE} & 0 & \omega_1 & -R_1^E-k_{EG} \end{pmatrix}\begin{pmatrix} I_x^G \\ I_y^G \\ I_z^G \\ I_x^E \\ I_y^E \\ I_z^E \end{pmatrix}$$

where $R_1^{G/E}$ is the longitudinal relaxation rate of the ground/excited state, $R_2^{G/E}$ is the transverse relaxation rate of the ground/excited state, $\omega_{G/E}$ is the offset of the applied $^{13}C$

spin lock with a strength of $\omega_1$ from the chemical shift ($\Omega_{G/E}$) of the ground/excited state, and $k_{GE}$ and $k_{EG}$ are forward and backward exchange rates as defined by $k_{GE} = p_E\, k_{ex}$ and $k_{EG} = p_G\, k_{ex}$. Here, $k_{ex} = k_{GE} + k_{EG}$ is the rate of exchange, $p_G$ and $p_E$ are populations of ground and excited states, respectively, and $\Omega_{obs} = p_G\, \Omega_G + p_E\, \Omega_E$ and $\Omega_E = \Omega_G + \Delta\omega$, where $\Delta\omega$ is the chemical shift difference between the ground and excited states. Ground state and excited state magnetizations at the beginning of the $T_{EX}$ period are along the effective spin-lock field as, $I_x^{G/E} = p_{G/E} \sin(\theta)$, $I_y^{G/E} = 0$, $I_z^{G/E} = p_{G/E} \cos(\theta)$, where $\theta = \arctan(\omega_1/\Omega)$ is the effective tilt angle. Fitting parameters are $R_1 = R_1^{G/E}$, $R_2 = R_2^{G/E}$, $\Delta\omega$, $k_{ex}$, and $p_E$, where we assume $R_1^G = R_1^E$ and $R_2^G = R_2^E$. Since the applied $^{13}C$ spin-lock powers are strong enough to decouple C-C couplings, the relaxation dispersion profiles of C1', C5, and C6 were analyzed the same as C2 and C8 profiles. For global fitting of dispersion profiles at individual pH condition, spin specific $R_1$, $R_2$, and $\Delta\omega$ were used, whereas $k_{ex}$ and $p_E$ were fit globally. For residues without conformational exchange, the two-state model was simplified to a one-state model by fixing all exchange parameters (rate of exchange $k_{ex}$ and population of excited state $p_E$) to 0. All profiles were fitted using an in-house OriginLab® program with a Levenberg-Marquardt algorithm.

**$^{15}N$ CEST measurements and data analysis.** $^{15}N$ CEST profiles were measured using a recently developed $^2J_{NH}$-based 2D $^1H$-$^{15}N$ HSQC CEST experiment pulse sequence that monitors longitudinal two-spin order ($2N_zH_z$) as described previously[42]. The $^{15}N$ carrier was set to 224.9 ppm with a spectral width of 6 ppm, and the $^{15}N$ offsets ranged between -6000 Hz and 3000 Hz with a spacing of 200 Hz, except for a of spacing of 50 Hz from -4200 to -3800 Hz and from 600 to 1000 Hz. With a relaxation period of $T_{EX}$ = 0.1 s, two $^{15}N$ $B_1$ fields ($\omega/2\pi$) of 52.1 Hz and 104.7 Hz were used at pH 8.04 and one $^{15}N$ $B_1$ field (ω/2π) of 52.1 Hz was used at pH 7.47 and 6.96. $^{15}$N spin-lock powers were calibrated according to the 1D approach by Guenneugues *et al.*[56] as previously described[51,57]. For all measurements, three spectra with $T_{EX}$ = 0 s were recorded for reference in data fitting and error estimation.

CEST profiles were analyzed as described previously[42]. Briefly, CEST profiles were obtained by normalizing peak intensity as a function of spin lock offset Ω to the peak intensity recorded at $T_{EX}$ = 0, where $\Omega = \omega_{rf} - \Omega_{obs}$ is the difference between the spin-lock carrier ($\omega_{rf}$) and the observed peak ($\Omega_{obs}$) frequencies. Measurement errors were estimated based on both triplicates performed at $T_{EX}$ = 0 s and the baseline of CEST profiles. Two-spin order $^{15}$N CEST profiles for A22-N1 at various pHs displaying conformational exchange were fit to a two-state exchange model between the ground (G) and excited (E) states based on the Bloch-McConnell equation[58] that describes magnetization evolution in a coupled two-spin $^{15}$N-$^{1}$H system[59,60],

$$\frac{d}{dt}\mathbf{v}^{G/E} = -\mathbf{R}^{G/E}\mathbf{v}^{G/E} = \begin{pmatrix} R_2^{G/E} & \omega_N^{G/E} & 0 & \eta_{xy}^{G/E} & \pi J_{NH}^{G/E} & 0 \\ -\omega_N^{G/E} & R_2^{G/E} & \omega_1 & -\pi J_{NH}^{G/E} & \eta_{xy}^{G/E} & 0 \\ 0 & -\omega_1 & R_1^{G/E} & 0 & 0 & \eta_z^{G/E} \\ \eta_{xy}^{G/E} & \pi J_{NH}^{G/E} & 0 & R_{2HN}^{G/E} & \omega_N^{G/E} & 0 \\ -\pi J_{NH}^{G/E} & \eta_{xy}^{G/E} & 0 & -\omega_N^{G/E} & R_{2HN}^{G/E} & \omega_1 \\ 0 & 0 & \eta_z^{G/E} & 0 & -\omega_1 & R_{1HN}^{G/E} \end{pmatrix} \begin{pmatrix} N_x^{G/E} \\ N_y^{G/E} \\ N_z^{G/E} \\ 2H_zN_x^{G/E} \\ 2H_zN_y^{G/E} \\ 2H_zN_z^{G/E} \end{pmatrix}$$

$$\frac{d}{dt}\boldsymbol{\sigma}(\mathrm{t}) = -\mathbf{L}\cdot\begin{bmatrix}\mathbf{v}^G \\ \mathbf{v}^E\end{bmatrix} = \left(\begin{bmatrix}\mathbf{R}^G & \mathbf{0}_6 \\ \mathbf{0}_6 & \mathbf{R}^E\end{bmatrix} + \begin{bmatrix}-k_{GE} & k_{EG} \\ k_{GE} & -k_{EG}\end{bmatrix} \otimes \mathbf{1}_6\right)\cdot\begin{bmatrix}\mathbf{v}^G \\ \mathbf{v}^E\end{bmatrix}$$

where $\mathbf{v}^{G/E}$ is the magnetization matrix, $\mathbf{R}^{G/E}$ is the relaxation matrix, $R_1^{G/E}$ is the $^{15}$N longitudinal relaxation rate, $R_2^{G/E}$ is the $^{15}$N transverse relaxation, $R_{1HN}^{G/E}$ is the $^{15}$N-$^{1}$H two-spin order relaxation rate, $R_{2HN}^{G/E}$ is the $^{15}$N antiphase relaxation rate, $\eta_z^{G/E}$ is the N-H dipolar-dipolar/nitrogen CSA cross-correlated relaxation between the $^{15}$N longitudinal and two-spin order elements, $\eta_{xy}^{G/E}$ is N-H dipolar-dipolar/nitrogen CSA cross-correlated relaxation between $^{15}$N transverse and antiphase magnetizations, $\omega_N^{G/E}$ is the offset of the applied $^{15}$N $B_1$ field with a strength of $\omega_1$, and $J_{NH}^{G/E}$ is the $^{15}$N-$^1$H scalar coupling for the ground (G) and excited (E) states, and $k_{GE}$ and $k_{EG}$ are forward and backward exchange rates as defined by $k_{GE} = p_E\ k_{ex}$ and $k_{EG} = p_G\ k_{ex}$. Here, $k_{ex} = k_{GE} + k_{EG}$ is the rate of exchange, $p_G$ and $p_E$ are populations of ground and excited states, respectively, and $\omega^G = \Omega_{obs}$ and $\omega^E = \omega^G + \Delta\omega$, where $\Delta\omega$ is the chemical shift difference between the ground and excited states. Ground state and excited state magnetizations at the beginning of the $T_{EX}$ period are the two-spin order ($2N_zH_z$) along Z and are set to be at populations of $p_G$ and $p_E$. The fitting parameters are $\Delta\omega$, $k_{ex}$, $p_E$, $R_2 = R_2^{G/E}$, $R_{1HN} = R_{1HN}^{G/E}$, and $R_{2HN} = R_2^{G/E} + R_{1HN}^{G/E} - R_1^{G/E}$ as described previously[42,61]. To simplify data fitting, $\eta_z^{G/E}$ and $\eta_{xy}^{G/E}$ were set to 0 as they have been shown not to affect the extracted $\Delta\omega$, $k_{ex}$, and $p_E$ values[42,62], and $R_1^{G/E}$ were set to 0, as the data does not constrain determination of $R_1$[42]. For residues without conformational exchange, the two-state model was simplified to a one-state model by fixing all exchange parameters (rate of exchange $k_{ex}$ and population of excited state $p_E$) to 0. All profiles were fitted using an in-house OriginLab® program with a Levenberg-Marquardt algorithm.

**Preparation of Dicer substrates.** Pre-miR-21 substrates were prepared by *in vitro* transcription, where a hammerhead ribozyme (HH) was fused to the 5'-end of pre-miR-21s to generate 5'-OH substrates with wild-type 5'-end nucleotide. Sequences of the pre-miR-21 substrates are as follows,

HH-Pre-miR-21 WT:

CUAAUACGACUCACUAUAGGAGCUACUGAUGAGGCCGAAAGGCCGAAACCCGAA
AGGGUCUAGCUUAUCAGACUGAUGUUGACUGUUGAAUCUCAUGGCAACACCAGU
CGAUGGGCUGUC

HH-Pre-miR-21 GC Mutant:

CUAAUACGACUCACUAUAGGAGCUACUGAUGAGGCCGAAAGGCCGAAACCCGAA
AGGGUCUAGCUUAUCAGACUGAUGUUGACCGUUGAAUCUCACGGCAACACCAGU
CGAUGGGCUGUC

HH-Pre-miR-21 G38U Mutant:

CUAAUACGACUCACUAUAGGAGCUACUGAUGAGGCCGAAAGGCCGAAACCCGAA
AGGGUCUAGCUUAUCAGACUGAUGUUGACUGUUGAAUCUCAUGUCAACACCAGU
CGAUGGGCUGUC

After *in vitro* transcription, samples were exchanged into $H_2O$ using Amicon filtration units (Millipore) with 10K Da MW cut-off membranes. The resulting substrates were subsequently subject to 5'-end $^{32}$P-labeling with T4 Polynucleotide Kinase (T4PNK) (New England Biolabs Inc.). 50 μl of RNA at 400 nM was incubated in a solution containing T4PNK buffer, 20 U T4PNK, and γ-$^{32}$P ATP at 35°C for 30 minutes before being gel purified by 20% denaturing polyacrylamide gel. The $^{32}$P-labeled RNAs were extracted from the gel by crushing and soaking in 250 mM NaCl in 1xTBE buffer for 24 hours before filtering and exchanging multiple times into $H_2O$ using an Amicon filtration unit (Millipore) with a 10K Da MW cut-off. $^{32}$P-labeled substrates were stored at -20°C prior to dicer processing assay.

**Dicer processing assay.** $^{32}$P-labeled pre-miR-21 substrates were heated to 95°C for 5 minutes and then placed on ice for 5 minutes to anneal the precursor hairpin. 4 μl substrate was mixed with 2 μl of 5x dicing buffer (12 mM HEPES, 1 M NaCl, and 0.02 mM EDTA at pH 7.8) and 2 μl of 25 mM $MgCl_2$. In a separate tube, concentrated Dicer enzyme (Genlantis, Inc.) was diluted in 1x dicing buffer (2.4 mM HEPES, 200 mM NaCl, 4 μM EDTA) and mixed with an equal volume of 10 mM ATP. 2 μl of ATP/Dicer mixture (10 mM ATP and 0.2U Dicer) was added to each tube of 8 μl substrate mixture and incubated at 35°C overnight. Reactions were quenched by adding 1.6 U proteinase K (New England Biolabs Inc.), 2 μl of 0.5 M EDTA and incubated for 45 minutes at 35°C. Samples were denatured in formamide with trace bromophenol blue and xylene cyanol, ran on a 15% denaturing polyacrylamide gel, visualized using Amersham Typhoon 5 Biomolecular Imager (GE Healthcare) and analyzed with Image Quant (GE Healthcare) using peak area and rubberbanding for background deletion. All assays were carried out in quadruplicate to estimate experimental errors.

**RNA secondary structure prediction.** All RNA secondary structures were predicted based on sequences using programs Mfold[63] and MC-Fold[45] using standard input options.

**Data availability.** The authors declare that the data supporting the findings of this study are available within the article and its Supplementary Information files, or are available upon reasonable request.

**Code availability.** The in-house OriginLab® scripts for data analyses are available upon request.

### SUPPLEMENTARY FIGURES

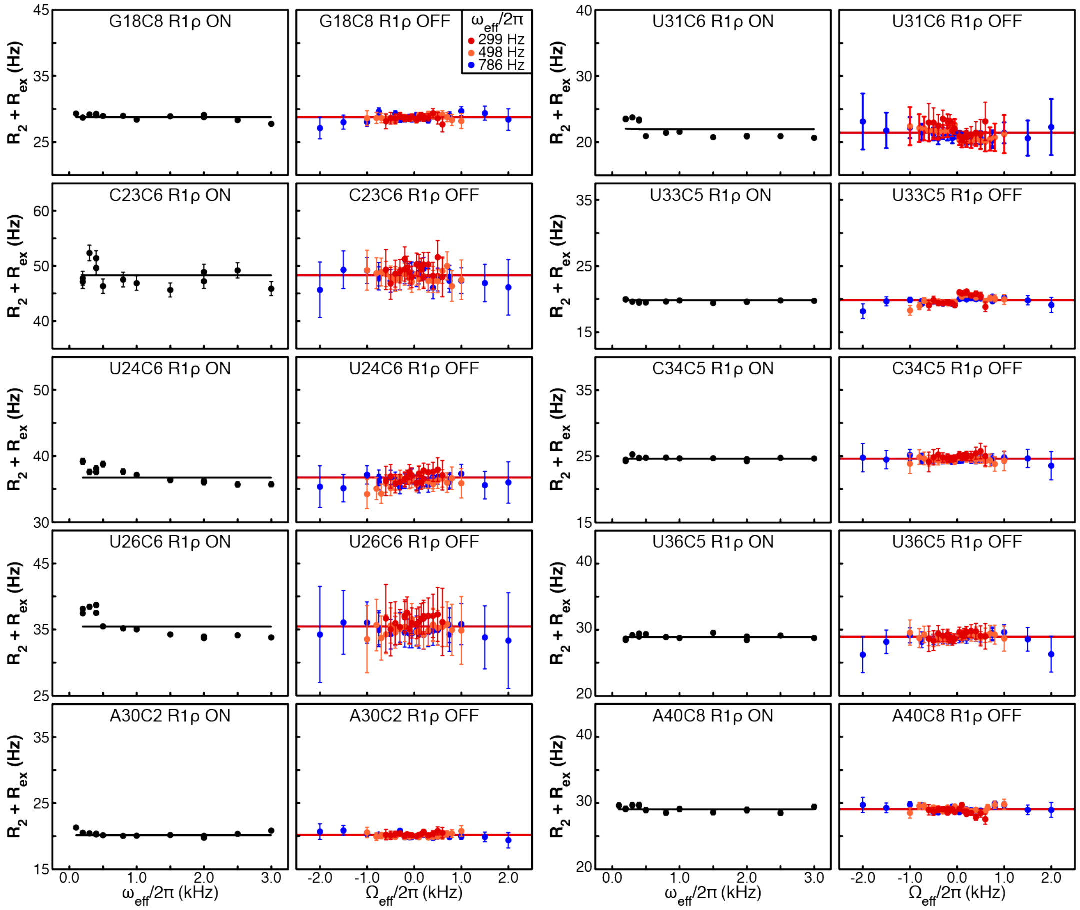

**Supplemental Figure 1** $^{13}C$ $R_{1\rho}$ RD profiles of preE-miR-21 residues without apparent chemical exchange. On- and off-resonance $^{13}C$ RD profiles depicting spin-lock power ($\omega_{eff}/2\pi$) and offset ($\Omega/2\pi$) dependence of $R_2 + R_{ex}$ measured at pH 6.45. Solid lines represent the best fits to a single-state model using the Bloch-McConnell equation. Error bars, experimental uncertainties (s.d.) estimated from mono-exponential fitting of $n$ = 3 independently measured peak intensities.

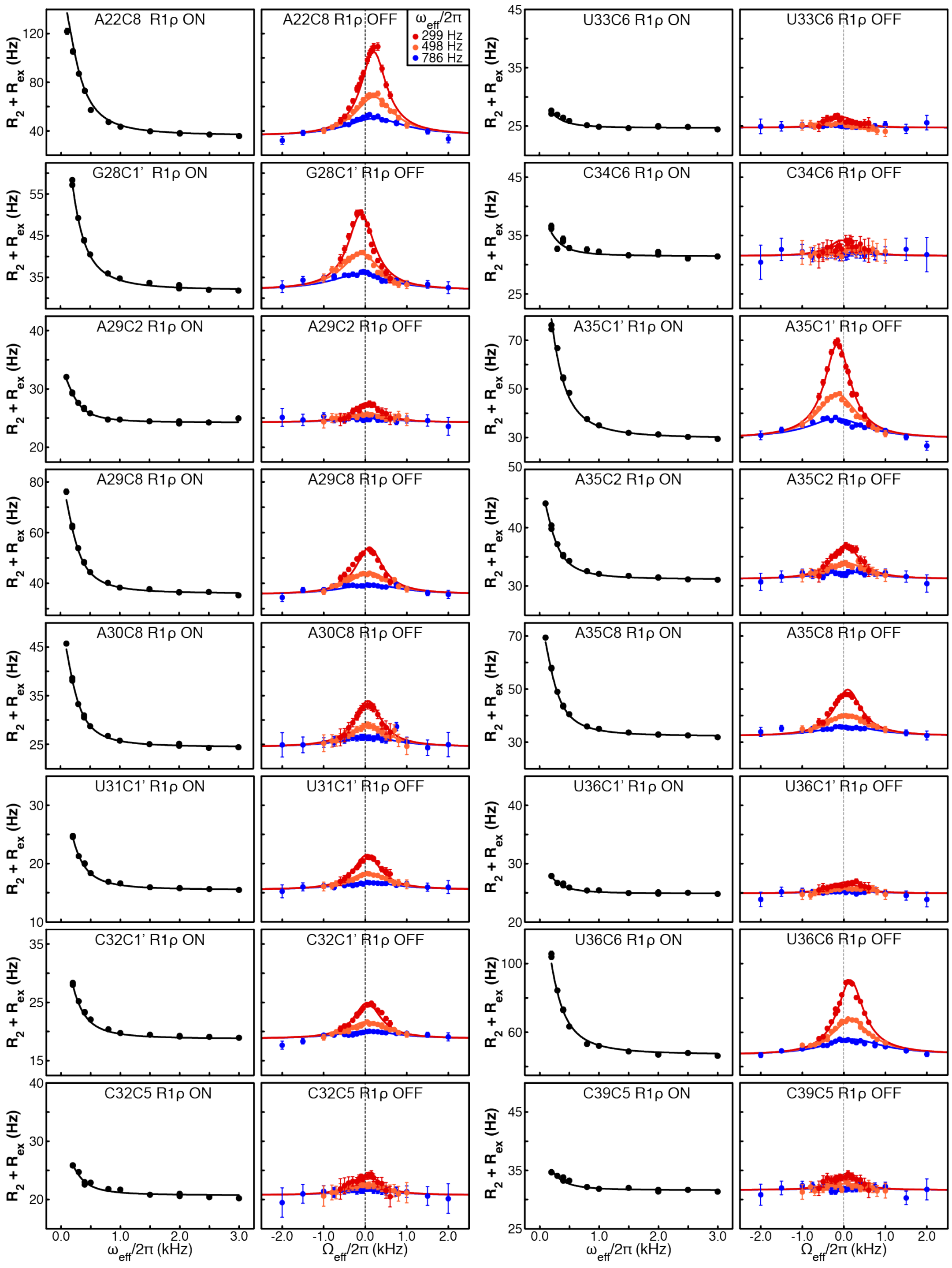


**Supplemental Figure 2** $^{13}$C $R_{1\rho}$ RD profiles of preE-miR-21 residues that undergo chemical exchange at pH 6.45. Solid lines represent the best fits to a global two-state exchange model ($k_{ex}$ = 1445 ± 17 s$^{-1}$ and $p_E$ = 15.2 ± 0.3%) using the Bloch-McConnell equation. Error bars, experimental uncertainties (s.d.) estimated from mono-exponential fitting of $n$ = 3 independently measured peak intensities.

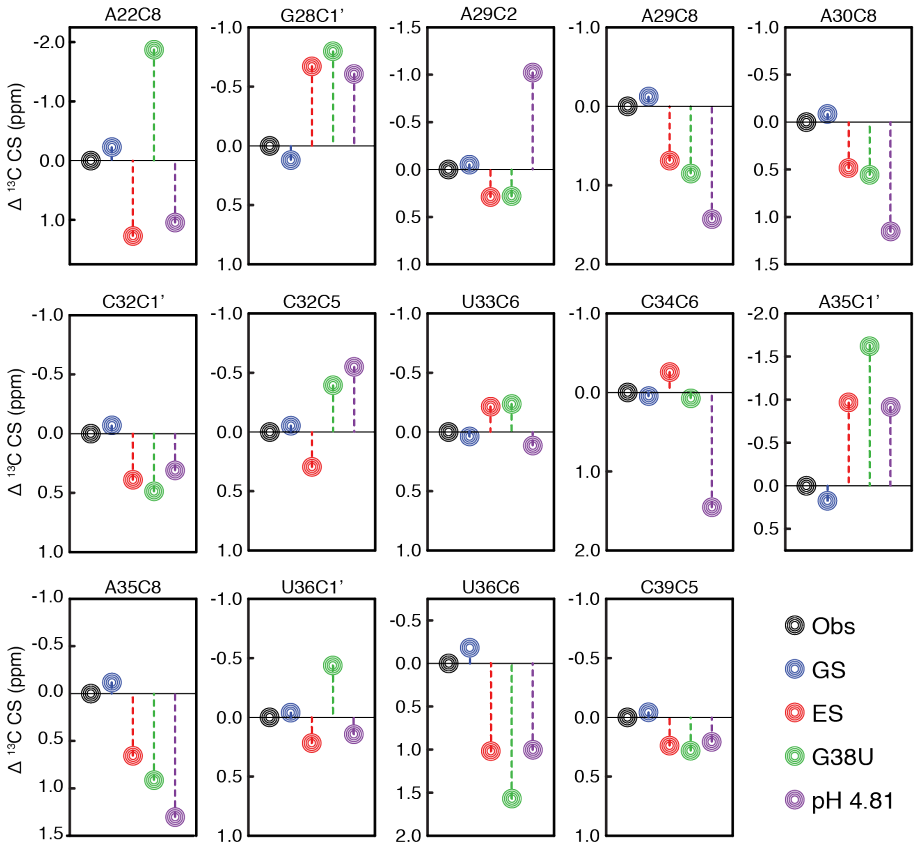


**Supplemental Figure 3** Comparison of GS, ES, mutant carbon chemical shifts. Shown are differences between the observed chemical shifts (black) and the chemical shifts of GS (blue) and ES (red) extracted from 13C R1ρ RD profiles at pH 6.45, G38U mutant at pH 6.45, and preE-miR-21 at pH 4.81. The apparent discrepancies between ES base carbon chemical shifts of A-C2/C8s and C-C5/C6s and their corresponding chemical shifts at pH 4.81 are likely due to intrinsic protonation at A-N1s and C-N3s at pH 4.81.

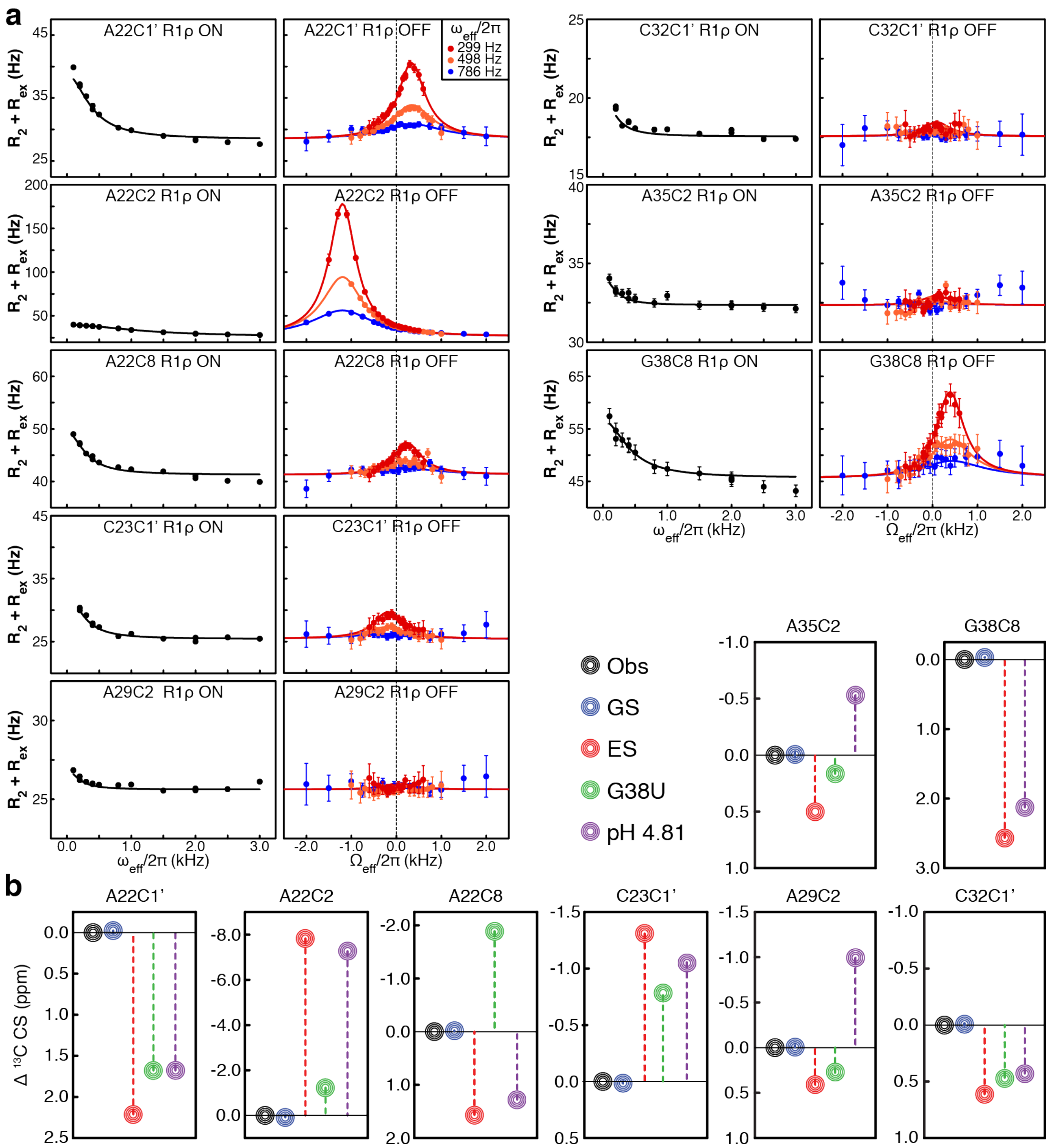


**Supplemental Figure 4** $^{13}$C $R_{1\rho}$ RD characterization of preE-miR-21 at pH 8.04. (**a**) On- and off-resonance $^{13}$C RD profiles of residues that are exchange-broadened at lower pH points, including A22-C2, G38-C8, and G38-C1'. Solid lines represent the best fits to a global two-state exchange ($k_{ex}$ = 1228 ± 51 s$^{-1}$ and $p_E$ = 1.1 ± 0.1%) using the Bloch-McConnell equation. (**b**) Comparison of GS, ES, mutant carbon chemical shifts. Shown are differences between observed chemical shifts (black) and chemical shifts of GS (blue) and ES (red) extracted from $^{13}$C $R_{1\rho}$ RD profiles at pH 8.04, G38U mutant at pH 6.45, and preE-miR-21 at pH 4.81. Error bars, experimental uncertainties (s.d.) estimated from mono-exponential fitting of $n$ = 3 independently measured peak intensities.

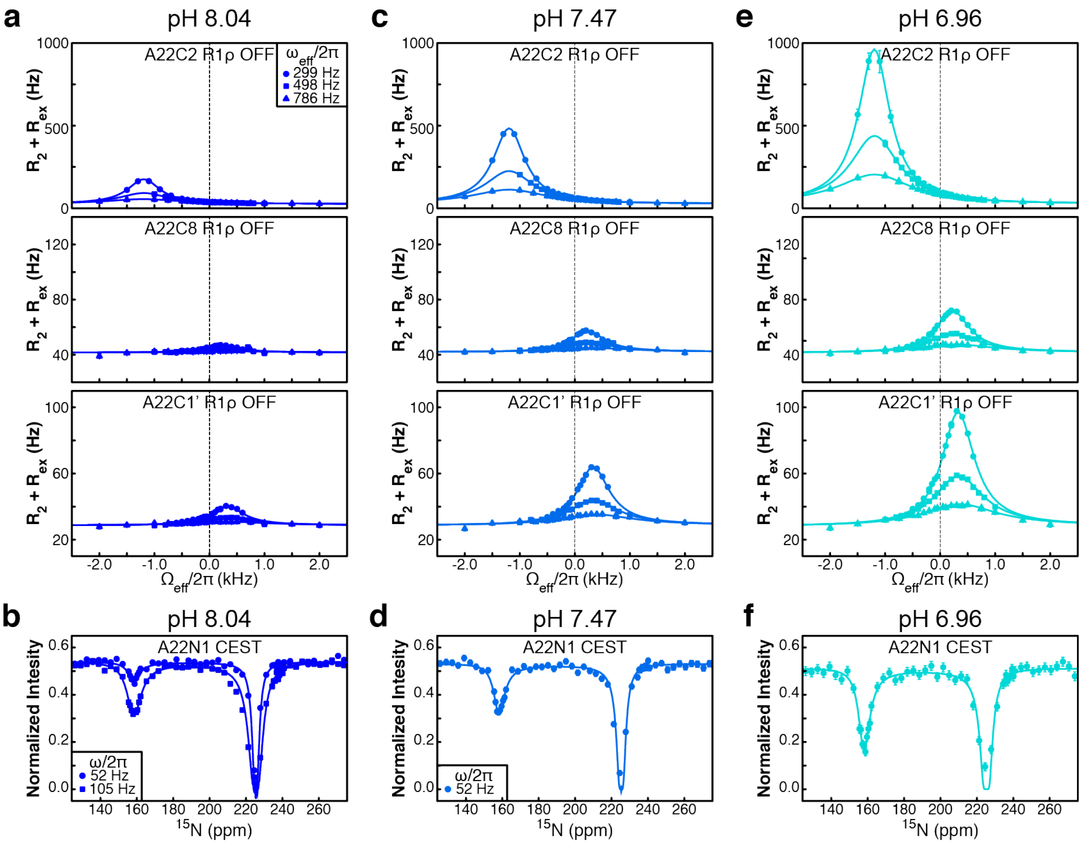


**Supplemental Figure 5** NMR RD characterization of pH-dependent chemical exchange of residue A22. Shown are $^{13}C$ $R_{1\rho}$ RD profiles of A22-C2/C8/C1' and $^{15}N$ CEST profile of A22-N1 at (**a-b**) pH 8.04, (**c-d**) pH 7.47, and (**e-f**) pH 6.96. Solid lines represent the best fits to a global two-state exchange model at each individual pH condition using the Bloch-McConnell equation, resulting in $p_E$ = 1.1 ± 0.1% at pH 8.04, $p_E$ = 3.4 ± 0.1% at pH 7.47, and $p_E$ = 6.4 ± 0.1% at pH 6.96. Error bars, experimental uncertainties (s.d.) estimated from $n$ = 3 independently measured peak intensities for CEST profiles and mono-exponential fitting of $n$ = 3 independently measured peak intensities for $R_{1\rho}$ RD profiles.

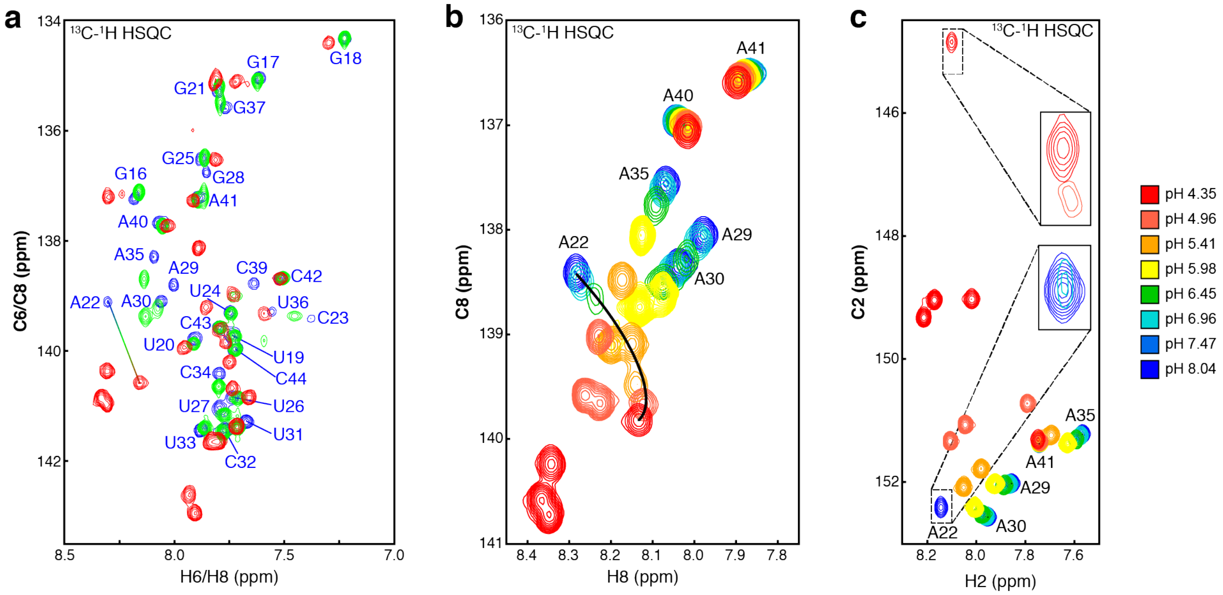


**Supplemental Figure 6** NMR characterization of pH-dependent changes of preE-miR-21. (**a**) $^{13}C$-$^{1}H$ HSQC spectra of base carbons (C6 and C8) of uniformly $^{13}C/^{15}N$-labeled preE-miR-21 at pHs 4.35, 6.45 and 8.04. (**b**) $^{13}C$-$^{1}H$ HSQC spectra of base carbons (C8) of adenine-specifically $^{13}C/^{15}N$-labeled preE-miR-21 with pH ranging from 4.35 to 8.04. (**c**) $^{13}C$-$^{1}H$ HSQC spectra of base carbons (C2) of adenine-specifically $^{13}C/^{15}N$-labeled preE-miR-21 with pH ranging from 4.35 to 8.04.

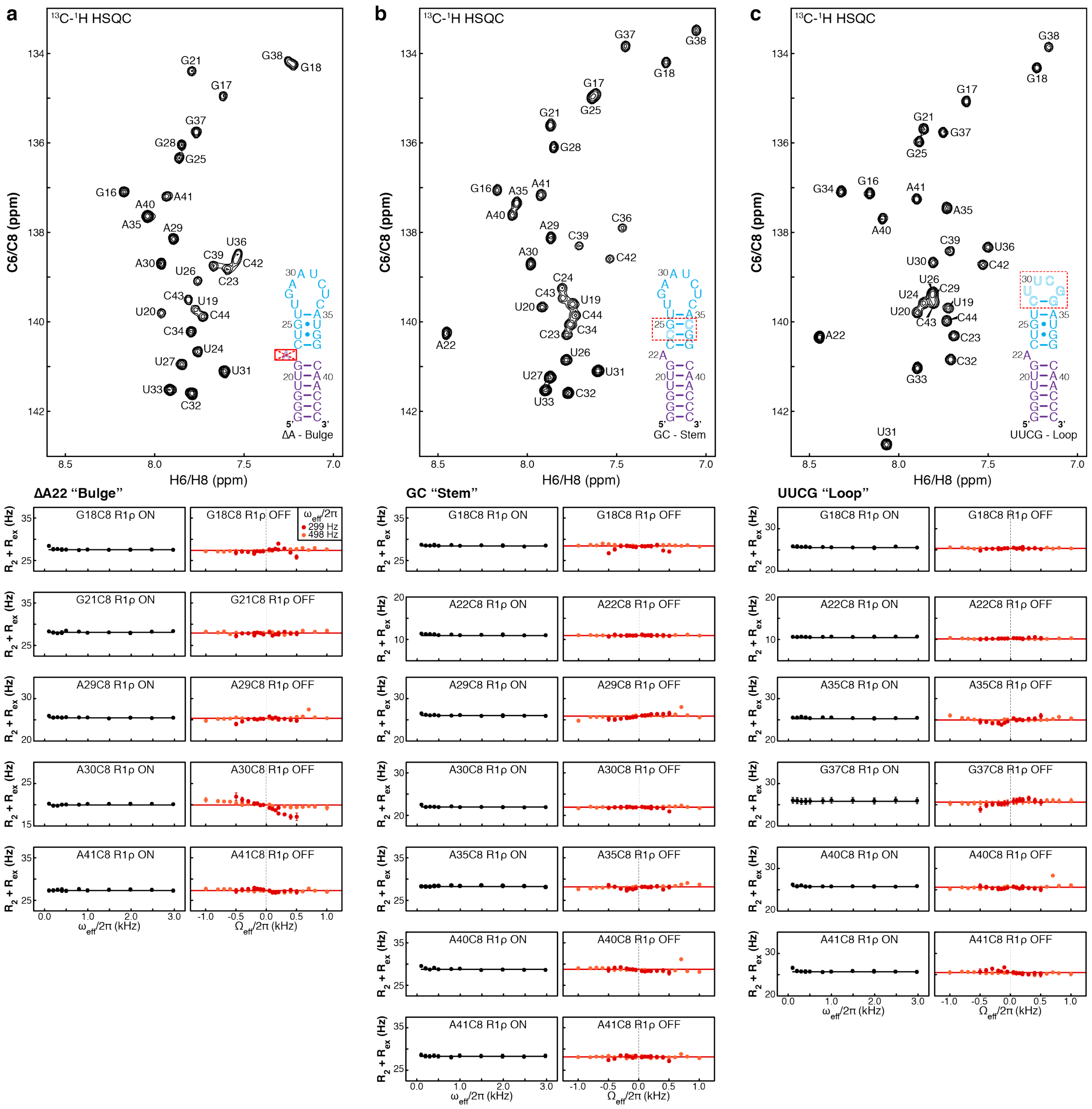

**Supplemental Figure 7** NMR characterization of preE-miR-21 mutants. Shown are $^{13}C$-$^{1}H$ HSQC spectra of base carbons (C6 and C8) and $^{13}C$ $R_{1\rho}$ RD profiles of uniformly $^{13}C/^{15}N$-labeled (**a**) ΔA22 "bulge", (**b**) GC "Stem", and (**c**) UUCG "Loop" mutants at pH 6.45. Solid lines represent the best fits to a single-state model using the Bloch-McConnell equation. Error bars, experimental uncertainties (s.d.) estimated from mono-exponential fitting of n = 3 independently measured peak intensities.

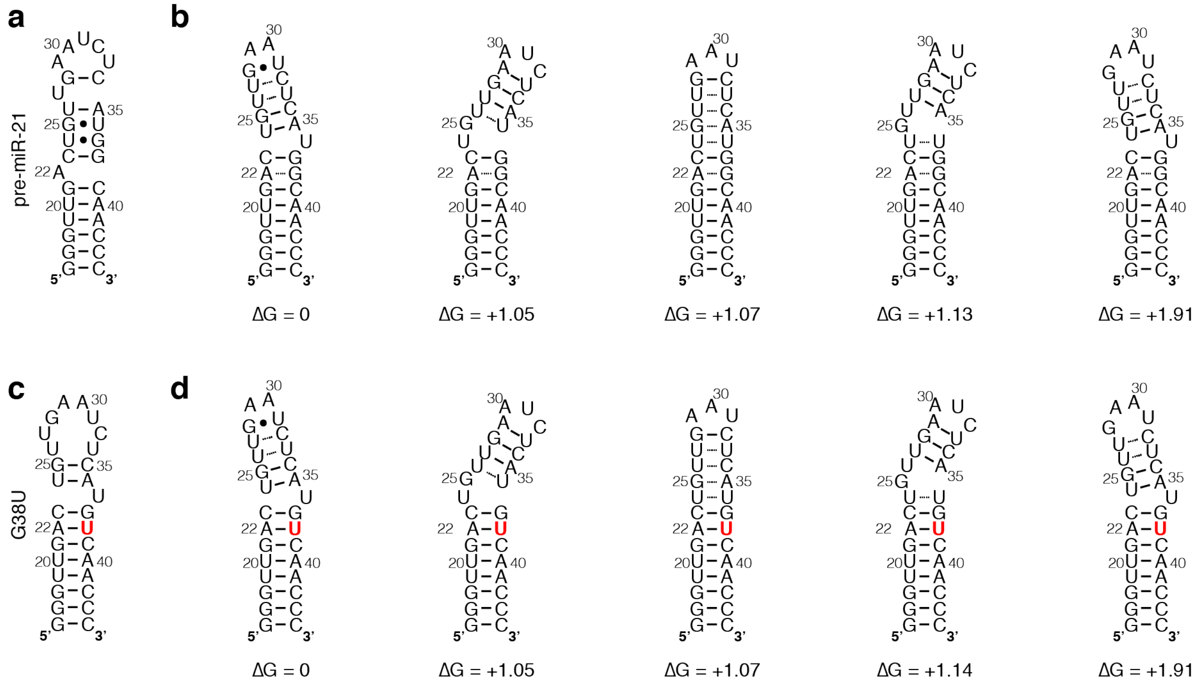


**Supplemental Figure 8** Secondary structure prediction of preE-miR-21 and G38U mutant by Mfold and MC-Fold. (**a**) M-fold predicted secondary structure of preE-miR-21, which contains G-U base pairs as observed in NMR data of the ground state. (**b**) MC-Fold predicted secondary structures of preE-miR-21. Shown are the top 5 lowest-energy structures, ranked with ΔG relative to the lowest predicted structure. The A22-G38 base pair is predicted in all MC-Fold structures. (**c**) M-fold predicted secondary structure of preE-miR-21 G38U mutant. (**d**) MC-Fold predicted secondary structures of preE-miR-21 G38U mutant. Shown are the top 5 lowest-energy structures, ranked with ΔG relative to the lowest predicted structure. Both M-fold and MC-Fold predict the same lower stem structure of preE-miR-21 G38U mutant, which is also consistent with NMR data.

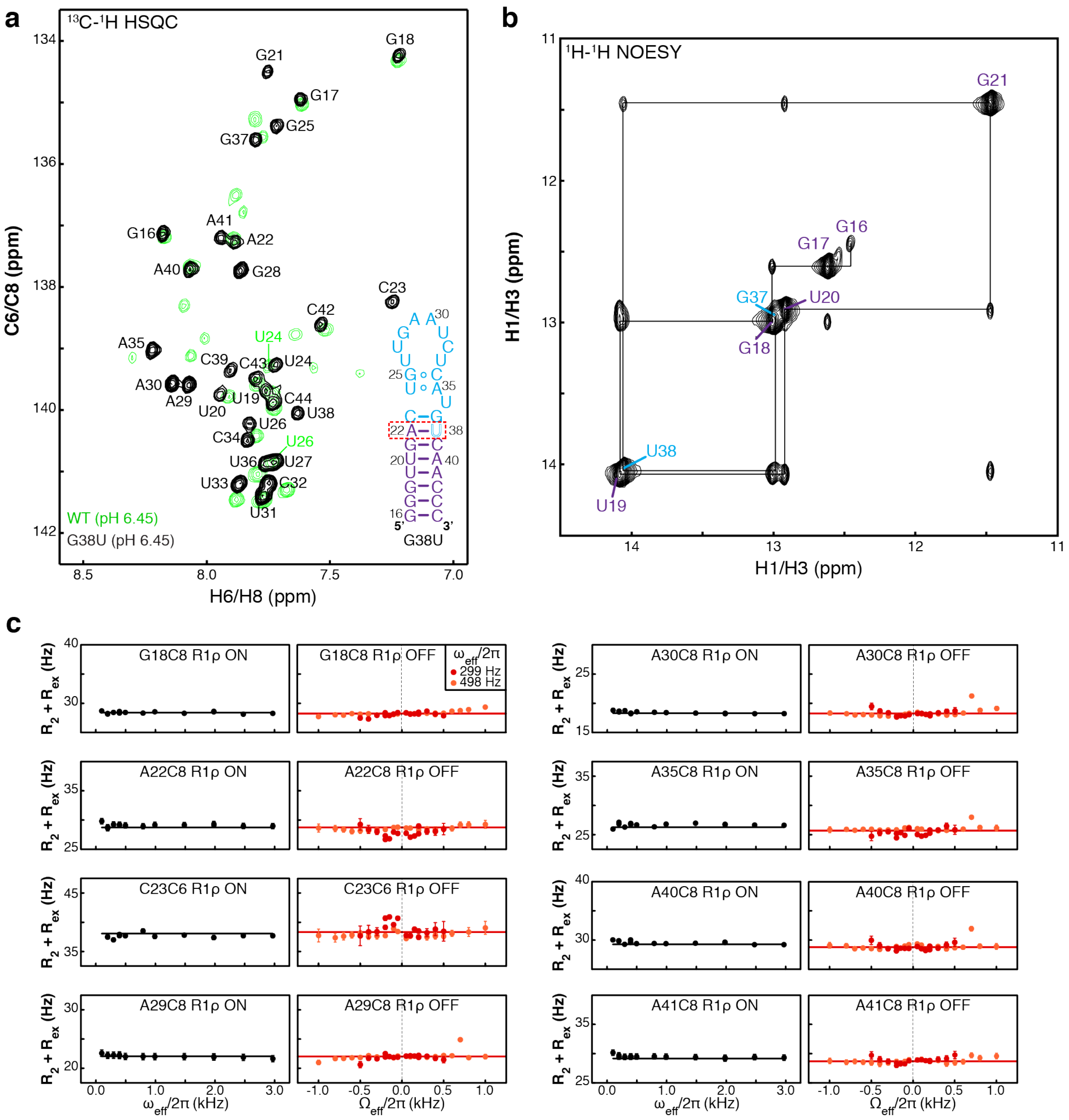

**Supplemental Figure 9** NMR characterization of preE-miR-21 excited-state mimic. Shown are (**a**) $^{13}C$-$^{1}H$ HSQC spectrum of base carbons (C6 and C8) (black) that overlays with $^{13}C$-$^{1}H$ HSQC spectrum of WT preE-miR-21 at pH 6.45 (green) (**b**) $^{1}H$-$^{1}H$ imino NOESY spectrum, and (**c**) $^{13}C$ $R_{1\rho}$ RD profiles of uniformly $^{13}C/^{15}N$-labeled G38U mutant at pH 6.45. Solid lines represent the best fits to a single-state model using the Bloch-McConnell equation. Error bars, experimental uncertainties (s.d.) estimated from mono-exponential fitting of $n$ = 3 independently measured peak intensities.
